# Adaptive Tracepoints for Pangenome Alignment Compression

**DOI:** 10.64898/2026.02.16.706236

**Authors:** Hasitha Kaushan, Santiago Marco-Sola, Erik Garrison, Pjotr Prins, Andrea Guarracino

## Abstract

**Motivation:** Storing millions of sequence alignments from large-scale genomic comparisons requires efficient compression methods. While fixed-size alignment encodings offer uniform spacing and bounded reconstruction cost, they cannot adapt to variable alignment complexity across sequences, missing compression opportunities in conserved regions.

**Results:** We present adaptive tracepoints, a complexity-aware alignment encoding that segments alignments using configurable complexity metrics (edit distance or diagonal distance) rather than fixed intervals. Segments are bounded by either the number of differences or the deviation from the main diagonal, adapting to local alignment characteristics. Reconstruction guarantees that alignments maintain identical or improved alignment scores. We validate the correctness of our method on simulated and real pangenomes with varying lengths and divergences. Diagonal-bounded tracepoints achieve 10.5-13.7*×* better compression than fixed-length encodings (*l*=100) on simulated long sequence alignments (100 Kb), while edit-bounded tracepoints provide a tunable trade-off between compression and reconstruction cost, approaching diagonal-bounded compression at higher thresholds with substantially lower memory and runtime. On real pangenomes (390M alignments), these methods compress alignments by 23-139*×* relative to uncompressed representations, with no score degradation and reconstruction time linear in alignment length.

**Availability:** Code and documentation are publicly available at https://github.com/AndreaGuarracino/tracepoints, https://github.com/AndreaGuarracino/tpa, and https://github.com/AndreaGuarracino/cigzip.

**Supplementary information:** Supplementary data are available at *Bioinformatics* online.

## 1 Introduction

Sequence alignment is fundamental to computational biology, enabling comparative genomics, variant detection, and evolutionary studies (Chao *et al*., 2022). However, the exponential growth of sequencing data poses major challenges for storing alignment information, particularly for large-scale genomic comparisons (Stephens *et al*., 2015; Greenfield *et al*., 2019). The CIGAR (Compact Idiosyncratic Gapped Alignment Report) string has emerged as the standard format for representing sequence alignments (Li *et al*., 2009). While CIGAR strings provide a complete representation of alignment operations, they require large storage, particularly for long-read sequencing and whole-genome comparisons. This storage burden has motivated the development of compressed alignment representations that balance space efficiency with the ability to reconstruct the original alignment.

For read-mapping applications, formats such as CRAM (Bonfield *et al*., 2022) use CIGAR strings to reconstruct query sequences from a reference, avoiding the need to store sequences directly. However, in pangenome comparisons and whole-genome alignments, both sequences (query and target) are primary data that are maintained independently. In these pairwise alignment scenarios, the challenge shifts to compressing the CIGAR strings themselves, as both sequences are already stored and available for alignment reconstruction.

The recent FastGA sequence aligner (Myers *et al*., 2025) adopted fixed-length tracepoints, originally proposed for long-read alignment (Myers, 2014), as a space-efficient alignment encoding. Rather than storing every alignment operation, fixed-length tracepoints record alignment endpoints at regular intervals (typically every 100 bases), enabling alignment reconstruction by re-aligning these smaller segments. This approach achieves compression by replacing detailed operation sequences with sparse positional information.

However, fixed-length segmentation has two practical limitations in certain scenarios. First, insertions and deletions (indels) that span segment boundaries must be split, leading to reconstruction artifacts where individual segments may produce different indel placements than the original alignment. To maintain biological correctness, indels should not be split across segments, which can limit the handling of large structural variations relevant to current genomic analyses (Amarasinghe *et al*., 2020). Second, fixed-length segmentation cannot adapt to natural variation in alignment complexity. A segment containing only matches in a conserved region uses the same representation as one filled with mismatches and indels in a divergent region, missing compression opportunities.

In this work, we present adaptive tracepoints, a complexity-aware alignment encoding that segments alignments based on local complexity rather than fixed lengths. We introduce two complexity metrics (i.e., edit distance and diagonal distance), each bounding segment size by a user-defined threshold, creating larger segments in conserved regions and smaller ones in divergent regions. Reconstruction via Wavefront Alignment (WFA) (Marco-Sola *et al*., 2020) guarantees identical or improved alignment scores, and we validate this on simulated and real pangenomes spanning a range of sequence lengths and divergence levels.

## 2 Background

### 2.1 Sequence Alignment and CIGAR Representation

Sequence alignment is a core operation in genome analysis and bioinformatics, used to detect similarities and differences that support evolutionary analysis, functional annotation, and structure prediction, among other applications. Formally, let query *q*_1,*⃛*,*n*_ and target *t*_1,*⃛*,*m*_ be two sequences of symbols of length *n* and *m*, respectively. Sequence alignment involves computing a transformation from sequence *q* into *t* using four basic operations: match, mismatch, insertion, and deletion. The optimal alignment is the sequence of operations that transforms *q* into *t* while minimizing a given cost function, such as the edit distance. For this discussion, let *e* be the sequence’s dissimilarity measured in edit operations (or edit count) and *ϵ* = *e/n* ≤ 1 the error rate. Without loss of generality, we assume that *m* ≤ *n*(1 + *ϵ*) = *θ*(*n*).

Classically, sequence alignment is solved using dynamic programming (DP) algorithms, such as variants of Needleman-Wunsch (Needleman and Wunsch, 1970) and Smith-Waterman (Smith and Waterman, 1981), which compute optimal alignments in quadratic time and space. Other proposals, such as OND (Myers, 1986) and WFA (Marco-Sola *et al*., 2020) algorithms, present output-sensitive formulations, reducing the running time to *O*(*ns*), where *s* is the optimal alignment score. Regardless of the algorithm, an alignment is usually represented as a sequence of edit operations (i.e., match=M, mismatch=X, insertion=I, and deletion=D) that transform one sequence into the other.

Alignments are usually stored in the standard CIGAR format, which encodes them as a compact string of run-length alignment operations, where each operation is preceded by a count indicating how many consecutive times it is repeated. The CIGAR encoding allows the alignment to be delivered in *O*(*n*) time by explicitly storing all alignment operations. However, this representation requires storing *O*(*n* + *e*) space in the worst case. In practice, representing alignments in CIGAR format can lead to considerable storage overhead in large-scale alignment experiments.

### 2.2 Tracepoints and Fixed-Length Sampling

Instead of explicitly storing every alignment operation, the tracepoint sampling strategy records only a sparse set of coordinates along the alignment path. Each tracepoint marks the position at which the alignment crosses a specified coordinate. Then, any two consecutive tracepoints, *u* = (*u*_*i*_, *u*_*j*_) and *v* = (*v*_*i*_, *v*_*j*_), define a subalignment interval of the first sequence *q*[*u*_*i*_, *v*_*i*_] against an interval *t*[*u*_*j*_, *v*_*j*_] of the second. The tracepoint approach provides a continuum of space-time trade-offs between recomputing an alignment from scratch and explicitly storing every edit operation as in a full CIGAR encoding.

The fixed-length tracepoint (FL-TP) sampling, proposed by Myers (Myers, 2014) and adopted for genome alignment by Myers *et al*. (2025), proposes to store a tracepoint every *l* symbols on one of the two sequences. Consider the sequence *q*, FL-TP partitions it into segments *q*[0, *l*), *q*[*l*, 2*l*), …, *q*[(*N* − 1) · *l, n*), each of *l* symbols (except perhaps the last), and stores no more than *N* = ⌈*n/l*⌉ tracepoints (first and last are implicit). Note that the partitioning over *q* is fixed, but that over *t* can be arbitrary, since insertions and deletions can occur within any alignment interval. Nevertheless, indels within an alignment interval cannot exceed *e*. In other words, the subalignment region defined by two consecutive tracepoints has bounded dimensions: its height is fixed to *l*, and its width cannot exceed *l*+*e*. Thus, the maximum size of any subalignment rectangle is *l* × (*l* + *e*).

As a result, the FL-TP sampling requires storing no more than *n/l* tracepoints and log(*l* + *e*) bits per tracepoint, assuming that each tracepoint’s coordinates are stored as offsets from the previous one. In this case, note that we only require storing the tracepoint coordinate that splits *t*, since the positions on *q* fall exactly on the predetermined boundaries 0, *l*, 2*l*, …, making them implicit. Thus, the overall space required is *O*(*n* log(*l* + *e*)*/l*) = *O*(*n* log(*e*)) in the worst case.

Given a sequence of tracepoints *T*, we can reconstruct the alignment *A* by aligning each interval between consecutive tracepoints and concatenating the resulting subalignments. Using a DP-based alignment algorithm for reconstruction, the time cost per subalignment is *O*(*l*(*l* +*e*)) in the worst case. Since there are *N* = ⌈*n/l*⌉ subproblems, the overall reconstruction cost becomes *O*(*nl* + *le*) = *O*(*n* + *e*) in the worst case. In contrast, an output-sensitive alignment algorithm reduces the reconstruction cost to *O*(*le* + (*N* − 1)*l*) = *O*(*n* + *le*) = *O*(*n* + *e*) assuming the worst case when the edit operations are clustered in a single subalignment. In practice, FastGA (Myers *et al*., 2025) also stores the per-segment edit count alongside the target advance, providing a local edit bound that can accelerate reconstruction.

As a consequence, the parameter *l* provides a tunable trade-off, as smaller values reduce reconstruction time but increase storage, while larger values save space at the cost of increased reconstruction time. The FL-TP sampling partitions the alignment uniformly, using the same tracepoint density regardless of how edit operations are distributed. As a result, it inefficiently oversamples conserved regions and incurs unnecessary storage overhead. We claim the FL-TP uses a uniform tracepoint density that does not adapt to local alignment complexity, leading to unnecessary storage overhead.

## 3 Methods

In this work, we propose a novel alignment compression strategy based on adaptive tracepoint sampling. In contrast to the FL-TP strategy, we use variable-length intervals that adapt to local alignment complexity, as measured by the alignment operations between tracepoints. To this end, we introduce two complementary methods: edit-bounded tracepoint (EB-TP) sampling and diagonal-bounded tracepoint (DB-TP) sampling. EB-TP determines tracepoint intervals by bounding the alignment operations (i.e., edit cost) accumulated between consecutive tracepoints, whereas DB-TP determines intervals based on the diagonal drift observed between tracepoints. These complementary sampling strategies offer distinct space-time trade-offs, depending on the alignment complexity profile and cost model used. In short, Table 1 summarizes the asymptotic storage costs and reconstruction time bounds for CIGAR, FL-TP, EB-TP, and DB-TP alignment representations. Note that, as usual in *O*(·) notation, constant factors (e.g., *l, δ*, and *b*) are suppressed.

**Table 1.**
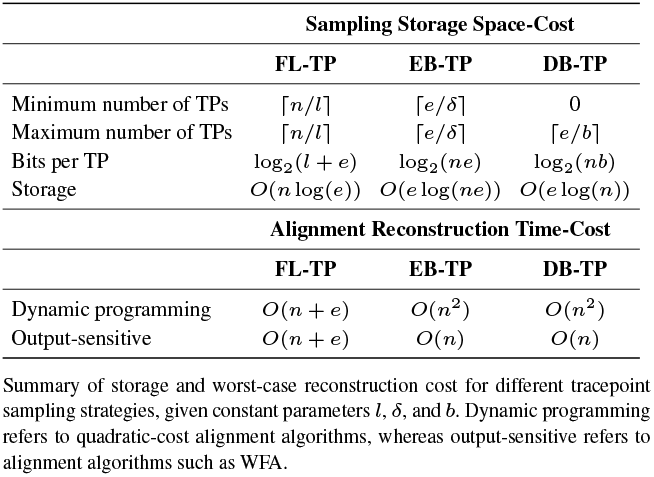
Tracepoint sampling strategies cost bounds.

| Sampling Storage Space-Cost |  |  |  |
| --- | --- | --- | --- |
|  | FL-TP | EB-TP | DB-TP |
| Minimum number of TPs | $\lceil n/l \rceil$ | $\lceil e/\delta \rceil$ | 0 |
| Maximum number of TPs | $\lceil n/l \rceil$ | $\lceil e/\delta \rceil$ | $\lceil e/b \rceil$ |
| Bits per TP | $\log_2(l + e)$ | $\log_2(ne)$ | $\log_2(nb)$ |
| Storage | $O(n \log(e))$ | $O(e \log(ne))$ | $O(e \log(n))$ |
| Alignment Reconstruction Time-Cost |  |  |  |
|  | FL-TP | EB-TP | DB-TP |
| Dynamic programming | $O(n + e)$ | $O(n^2)$ | $O(n^2)$ |
| Output-sensitive | $O(n + e)$ | $O(n)$ | $O(n)$ |

Summary of storage and worst-case reconstruction cost for different tracepoint sampling strategies, given constant parameters $l$ , $\delta$ , and $b$ . Dynamic programming refers to quadratic-cost alignment algorithms, whereas output-sensitive refers to alignment algorithms such as WFA.

### 3.1 Edit-Bounded Tracepoint Sampling

The edit-bounded tracepoint (EB-TP) sampling strategy partitions the alignment by content into intervals of at most *δ* edit operations. By adapting to the local edit rate, this strategy yields smaller and more balanced subproblems than fixed-length partitioning.

The EB-TP sampling requires storing *N* = ⌈*e/δ*⌉ tracepoints and 2 log_2_(*n*) bits per tracepoint, in the worst case, yielding segments of variable length in both axes. Using this strategy, the subalignment region between two consecutive tracepoints cannot be more skewed than an *n* × (*n* ± *e*) rectangle. Therefore, the second coordinate can be encoded as its difference from the first, requiring no more than (log_2_(*n*) +log_2_(*e*)) bits per tracepoint. In the worst case, the overall space required is *O*((*e/δ*) · log(*ne*)) = *O*(*e* log(*ne*)).

In practice, the adaptive EB-TP sampling is highly efficient when the sequences are similar. If the alignment contains few edit operations, the number of tracepoints collapses to a small constant because each new tracepoint is created only after every *δ* edits. When *e* ∼ *δ*, the sampling requires little space, far smaller than either fixed-length tracepoints or full CIGAR representations.

Regarding the alignment reconstruction, given an array of EB-TP sampled tracepoints, the full alignment can be recovered by recomputing the *N* partial alignments and concatenating them. Under the edit distance, if each subalignment between consecutive tracepoints is recomputed optimally, then the reconstructed alignment *Â* has total cost less than or equal to that of *A*. In particular, if *A* is not optimal, the reconstructed alignment cannot be worse and may strictly improve its overall alignment cost (Lemma S1.1 in Supplementary).

Regarding reconstruction cost, a quadratic DP-based algorithm requires *O*(*n*^2^) time to reconstruct the original alignment in the worst case. In contrast, an output-sensitive alignment algorithm would take *O*(*nδ*) = *O*(*n*) time in the worst case.

### 3.2 Diagonal-Bounded Tracepoint Sampling

Similarly, we propose a diagonal-aware strategy that samples tracepoints only when the alignment deviates significantly (*b* units) from its current diagonal. The key idea is to exploit the fact that high-similarity alignments remain close to a dominant diagonal and, therefore, tracepoints should be created only when the alignment path drifts too far from this diagonal, indicating a substantial local shift.

For that, the diagonal-bounded tracepoint (DB-TP) sampling strategy follows the alignment path and maintains the diagonal of the most recently sampled tracepoint as the reference diagonal. As it advances along the alignment, it monitors the deviation from the diagonal of the last sampled tracepoint. If the drift remains within a band of width 2*b* (i.e., *b* positions to the right and *b* to the left), no tracepoint is emitted.

The DB-TP sampling requires storing at most ⌈*e/b*⌉ tracepoints and (log_2_(*n*) + log_2_(*b*)) bits per tracepoint. In this case, the subalignment region between two consecutive tracepoints cannot be more skewed than an *n* × (*n* ± *b*) rectangle. In the worst case, the overall space required is *O*((*e/b*) · (log(*nb*))) = *O*(*e* log(*n*)).

As with the previous strategy, recomputing each subalignment optimally between consecutive DB-TP tracepoints guarantees a globally optimal alignment under edit distance (Lemma S1.1). In this case, a quadratic DP-based algorithm would require *O*(*n*^2^) time to reconstruct the original alignment. In contrast, an output-sensitive alignment algorithm requires *O*(*nb*) = *O*(*n*) in the worst case when the errors are clustered within a single subalignment.

### 3.3 Generalization to Affine-Gap and Atomic Gaps

We have described tracepoint sampling and alignment reconstruction under the unit-cost edit distance model. However, most bioinformatics applications rely on gap-affine cost models that better capture biological gap structure. The affine-gap model (Smith and Waterman, 1981; Gotoh, 1982) distinguishes between the cost of opening a gap and the cost of extending it, reflecting the fact that long insertions or deletions often arise from a single biological event rather than from multiple independent events. This way, the affine scoring favors contiguous gaps over fragmented ones, producing alignments that are more biologically meaningful.

However, extending tracepoint-based reconstruction to the affine-gap model introduces an important caveat. If tracepoints are allowed to fall within gaps, gaps may be split across multiple subalignments. During reconstruction, these gap fragments are treated independently and may incur additional gap-opening penalties. As a result, even when each subalignment between tracepoints is locally optimal, their concatenation is not guaranteed to produce a globally optimal alignment under the affine-gap scoring model.

To address this limitation, we enforce atomic gaps during tracepoint sampling. That is, tracepoints are never placed inside gaps, ensuring that each gap is treated as an indivisible unit and that the correct gap state is preserved at subalignment boundaries. Under this constraint, independently recomputing optimal subalignments between consecutive tracepoints and concatenating them yields a globally optimal alignment even under affine-gap scoring (Lemma S1.2).

### 3.4 Exploiting Local Edit-Bounds

Tracepoints delimit subproblems along the alignment, but they do not convey any information about the difficulty of reconstructing each segment. In particular, the FL-TP sampling ignores the distribution of edit operations along the alignment. In contrast, EB-TP and DB-TP place tracepoints based on the accumulated alignment cost rather than on sequence length. This yields variable-length intervals whose complexity is explicitly bounded by the number of edit operations they contain.

We propose exploiting the number of edit operations contained in the corresponding subalignment interval. This additional annotation provides a local edit-bound for each subalignment, offering an explicit upper bound on its reconstruction cost. The key idea is to employ banded alignment techniques during reconstruction. By knowing the number of edit operations in each subalignment, we can explicitly bound the width of the diagonal band required to contain the optimal path for that segment. As a result, reconstruction no longer needs to explore the whole region, but only a narrow band whose width is proportional to the local edit bound.

In the supplementary material (Section S3), we demonstrate that local edit-bounds directly translate into bounds on the diagonal band required for reconstruction. Specifically, Lemmas S3.1 and S3.2 show that, under both edit-distance and affine-gap, the optimal subalignment between two consecutive tracepoints is guaranteed to lie within a diagonal band whose width is determined by the number of edits or by the alignment score of that subalignment. These results motivate the use of banded alignment reconstruction between tracepoints, as the optimal path cannot drift arbitrarily far from the central diagonal once local edit bounds are fixed.

From a theoretical perspective, these bounds do not change the worst-case asymptotic complexity of output-sensitive reconstruction algorithms. However, in practice, local edit-bounds enable reconstruction to focus computation on the relevant regions, reducing unnecessary computation. This results in a consistent reduction in reconstruction time without compromising alignment optimality.

### 3.5 Tracepoints Alignment Format

To enable efficient storage and rapid random access, we introduce the TPA (TracePoint Alignment) format, a binary file format that applies compression strategies to tracepoints. A file-level header records the segmentation metric, its threshold parameter (*δ* for EB-TP, *b* for DB-TP, or the tracespacing *l* for FL-TP), and the alignment scoring scheme (edit distance, gap-affine, or dual gap-affine). Each alignment record stores per-segment value pairs: query and target advances for EB-TP and DB-TP, or, as in FastGA (Myers *et al*., 2025), edit distance and target advance for FL-TP, whose fixed query advance is already in the header. TPA decomposes each record into two independent streams whose encoding strategies are determined by the segmentation metric, exploiting metric-specific statistical regularities in the data (Supplementary Section 4). TPA files are paired with an index that maps each alignment record to its byte offset, enabling *O*(1) random access.

## 4 Results

We implement adaptive tracepoints in Rust, interfacing with WFA2-lib (Marco-Sola *et al*., 2020, 2023) for alignment reconstruction. We also provide cigzip, a complementary tool for CIGAR compression used in our evaluation. We evaluate our encoding on both simulated and real pangenomes covering different evolutionary distances. All executions were performed with 96 threads on a node running Debian GNU/Linux 12, equipped with an AMD EPYC 9274F CPU (2 Sockets x 24 Cores, 96 threads in total) and 755 GB of RAM.

### 4.1 Evaluation on Simulated Data

For each combination of sequence length (100 bp, 1 Kb, 10 Kb, 100 Kb) and divergence rate (1%, 5%, 10%, and 20%), we generated *N* =10,000 random sequence pairs and performed optimal alignments using the WFA algorithm with edit distance (unit costs for mismatch, insertion, and deletion).

Figure 2 shows compression ratios compared to uncompressed PAF files across alignment lengths and error rates, using BGZIP-compressed PAF (Bonfield *et al*., 2021) as a baseline because, like TPA, it supports random access to individual records. For short sequences (100 bp), BGZIP achieves the best compression (0.11-0.21 ratio across error rates) due to efficient entropy coding of short CIGAR strings. As sequence length increases, adaptive tracepoint methods outperform both BGZIP and fixed-length tracepoints (FL-TP, *l* = 100). At 100 Kb, DB-TP TPA (*b* = 32) achieves 10.5-13.7× better compression than FL-TP and 27-132× better than BGZIP across error rates. EB-TP TPA (*δ* = 32) achieves 1.1-5.4× improvement over FL-TP at 100 Kb, with larger gains at lower error rates. Compression ratios improve with divergence for all tracepoint methods because CIGAR size scales linearly with the total number of edit operations, whereas tracepoints encode each segment (regardless of how many operations it contains) with exactly two values, yielding total storage that grows sub-linearly with error count (Table 1).

**Fig. 1:**
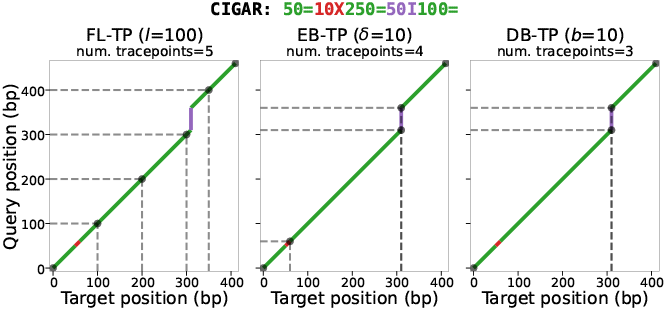
Visualization of a toy alignment under three tracepoint sampling strategies. Green indicates matches, red mismatches, and purple insertions. FL-TP produces fixed-length segments regardless of local complexity, while EB-TP and DB-TP adapt segment boundaries to alignment content.

**Fig. 2:**
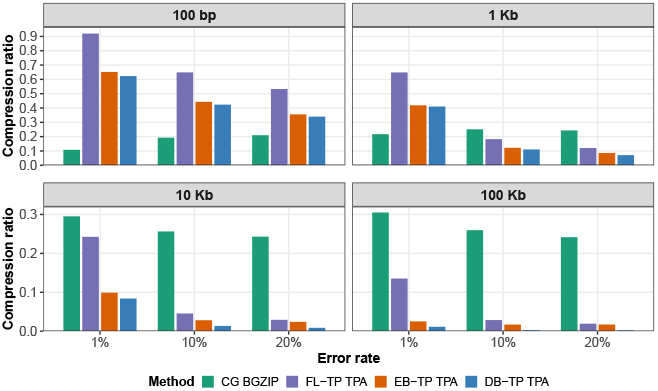
Compression ratios (compressed file size /uncompressed PAF file size) on simulated alignments (*N* =10,000 pairs per condition). Lower values indicate better compression. CG BGZIP = BGZIP-compressed PAF; FL-TP = fixed-length tracepoints (*l* = 100); EB-TP = edit-bounded tracepoints (*δ* = 32); DB-TP = diagonal-bounded tracepoints (*b* = 32). All tracepoint methods in TPA format.

DB-TP consistently outperforms EB-TP, with the advantage increasing with sequence length and error rate. This reflects the theoretical bounds in Table 1. The effect of varying the threshold (*δ* for EB-TP, *b* for DB-TP) across {32, 64, 128, 256, 512, 1024} is shown in Supplementary Figure S1.

Figure 3 compares reconstruction cost (TPA → PAF) against computing alignments from scratch and BGZIP decompression. BGZIP decompression is the fastest method at 100 Kb with negligible memory, but requires 12-132× more storage than adaptive tracepoint formats at 100 Kb. All tracepoint methods reconstruct individual alignments in sub-millisecond time at 100 Kb, up to 117× faster than re-alignment at high divergence. Adaptive methods generate fewer tracepoints than fixed-length encoding (Supplementary Figure S2). At 100 Kb, FL-TP produces ∼10M tracepoints regardless of divergence, while DB-TP produces only 130K at 10% divergence (77× fewer) and 20K at 1% divergence (∼500× fewer). EB-TP produces ∼2.9M tracepoints at 10% divergence, intermediate between FL-TP and DB-TP. The number of tracepoints directly affects decompression overhead. Decompression of the compressed TPA files accounts for at most 3% of total reconstruction time for DB-TP across all simulated conditions, confirming that its computational cost is dominated by segment re-alignment rather than I/O. In contrast, FL-TP’s ∼10M tracepoints at 100 Kb cause decompression to account for up to 71% of total time, and EB-TP’s intermediate tracepoint counts lead to decompression fractions of up to 26% at 100 Kb.

**Fig. 3:**
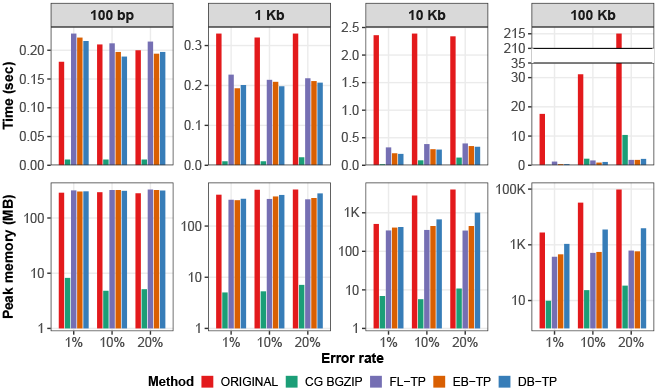
Reconstruction cost comparison. Top: decoding runtime; Bottom: peak memory usage. ORIGINAL = aligning sequences from scratch using WFA; CG BGZIP = BGZIP decompression to PAF; FL-TP, EB-TP, and DB-TP = TPA → PAF reconstruction for fixed-length (*l* = 100), edit-bounded (*δ* = 32), and diagonal-bounded (*b* = 32) tracepoints, respectively.

All reconstructions produced CIGAR strings (and, consequently, alignment scores) identical to those of the original input alignments, confirming our theoretical guarantee for optimal input alignments.

### 4.2 Evaluation on Real Data

Real data comprises two sets of heuristic pangenome alignments generated with WFMASH (Guarracino *et al*., 2021) with dual affine-gap scoring (mismatch 5, gap-open1 8, gap-ext1 2, gap-open2 24, gap-ext2 1):

- **Human pangenome:** 389,234,668 alignments in 466 PAF files (one per target haplotype) from 25,221 random haplotype pairs sampled from the Human Pangenome Reference Consortium Phase 2 pangenome (HPRCv2, 466 haplotypes). Total alignment length: 76.7 Tb; mean alignment length: 197 Kb (max: 30.1 Mb); mean alignment identity: 98.3%.
- **Primate pangenome:** 566,038 alignments in 16 PAF files (one per target haplotype) from inter-species all-versus-all comparisons of 16 haplotypes from the T2T ape genomes (Yoo *et al*., 2025): 12 haplotypes across 6 species (chimpanzee, bonobo, gorilla, Bornean and Sumatran orangutans, siamang) plus 4 human references (CHM13, GRCh38, HG002 maternal/paternal). Total alignment length: 683 Gb; mean alignment length: 1.21 Mb (max: 170 Mb); mean alignment identity: 91.2%.

Table 2 summarizes compression performance. DB-TP (*b*=32) achieves 0.025× compression ratio on the human pangenome and 0.007× on the primate pangenome. EB-TP ranges from 0.043× (*δ*=32) to 0.025× (*δ*=128) on human data, and from 0.025× to 0.009× on primate data. Alignment score distributions for both pangenomes are shown in Supplementary Figure S3.

**Table 2.** Performance comparison across two pangenomes. For BGZIP PAF, decompression time and memory refer to `bgzip -d` decompression; for tracepoint methods, they refer to full TPA→PAF reconstruction (decompression + segment re-alignment). Score improvement indicates the percentage of reconstructions that achieve better alignment scores than the input. All remaining reconstructions achieve identical scores; no reconstruction ever produces a worse score.

| Method | Human pangenome |  |  |  |  | Primate pangenome |  |  |  |  |
| --- | --- | --- | --- | --- | --- | --- | --- | --- | --- | --- |
|  | Storage (GiB) | Compr. ratio | Reconstr. time (h) | Reconstr. mem. (GiB) | Score improved | Storage (GiB) | Compr. ratio | Reconstr. time (h) | Reconstr. mem. (GiB) | Score improved |
| Uncompressed PAF | 940.10 | 1.000× | – | – | 0.00% | 78.90 | 1.000× | – | – | 0.00% |
| BGZIP PAF (level 9) | 296.51 | 0.315× | 0.14 | 0.08 | 0.00% | 22.01 | 0.279× | 0.01 | 0.04 | 0.00% |
| EB-TP TPA ($\delta=32$) | 40.15 | 0.043× | 4.38 | 15.03 | 0.15% | 2.00 | 0.025× | 0.18 | 17.30 | 68.34% |
| EB-TP TPA ($\delta=64$) | 29.31 | 0.031× | 4.64 | 15.59 | 0.64% | 1.15 | 0.015× | 0.19 | 17.74 | 76.58% |
| EB-TP TPA ($\delta=128$) | 23.27 | 0.025× | 5.28 | 17.92 | 3.15% | 0.70 | 0.009× | 0.25 | 19.16 | 80.07% |
| DB-TP TPA ($b=32$) | 23.63 | 0.025× | 10.78 | 65.06 | 0.54% | 0.57 | 0.007× | 4.47 | 247.75 | 75.66% |

For human-human alignments, DB-TP generates 4.7 billion tracepoints with a mean segment length of 16.3 Kb. EB-TP generates 11.0 billion (*δ*=32), 6.5 billion (*δ*=64), and 4.1 billion (*δ*=128) tracepoints, with mean segment lengths of 7.0, 11.8, and 18.9 Kb, respectively. That is, 2.3× (*δ*=32) to 0.9× (*δ*=128) the number of DB-TP tracepoints. For primate alignments, DB-TP produces 208M tracepoints (3.3 Kb segments), while EB-TP ranges from 885M (*δ*=32; 772 bp segments) to 266M (*δ*=128; 2.6 Kb segments), reflecting how diagonal-bounded segmentation exploits conserved regions. On the human pangenome, 99.46% of DB-TP reconstructions achieved identical scores to input alignments, compared to 99.85% (*δ*=32), 99.36% (*δ*=64), and 96.85% (*δ*=128) for EB-TP. The remaining reconstructions showed improved scores, with zero degradations in all cases: 0.54% (DB-TP) and 0.15%-3.15% (EB-TP, *δ*=32-128). On the primate pangenome, improvement rates were substantially higher: 75.66% (DB-TP) and 68.34%-80.07% (EB-TP, *δ*=32-128) of reconstructions improved upon the input alignments, again with zero degradations. This confirms our theoretical guarantee that reconstruction never produces worse alignments than the input, while demonstrating that exact WFA reconstruction often finds better paths than heuristic aligners, particularly for divergent sequences. Larger *δ* values impose fewer intermediate anchor points, giving exact WFA reconstruction more freedom to find optimal paths that improve upon the heuristic aligner’s original alignment.

On the human-human dataset, DB-TP reconstruction averages 83 seconds per file (10.8 hours total) with 65 GiB peak memory, while EB-TP averages 34-41 seconds (*δ*=32-128; 4.4-5.3 hours total) with 15-18 GiB peak memory. On the primate dataset, this gap widens: DB-TP averages 1,005 seconds per file (4.5 hours total) with 248 GiB peak memory, while EB-TP averages 40-57 seconds (*δ*=32-128; 0.2-0.3 hours total) with 17-19 GiB peak memory. That is, a 18-25× speed difference and 13-14× memory difference (Supplementary Figure S4). For comparison, BGZIP decompression completes in 0.14 hours (human) and 0.01 hours (primate) with *<* 1 GiB peak memory (Table 2), but requires 7-39× more storage than adaptive tracepoint formats. It is important to note that, according to our performance profiling results, up to 45% of CPU execution cycles are spent retrieving FASTA sequences from SSD for reconstruction. Also note that DB-TP’s higher reconstruction cost reflects its larger segment sizes: fewer but larger segments require more resources for WFA alignment within each segment. At *δ*=128, EB-TP achieves compression comparable to DB-TP on human data (0.025× vs 0.025×) while remaining 2-18× faster and requiring 4-13× less memory, making it competitive for workflows that balance storage and reconstruction cost. Decompression fractions follow the same pattern as simulations: DB-TP decompression accounts for under 2% of total reconstruction time across both pangenomes, while EB-TP decompression ranges from 6-10% (*δ*=32) to 2-3% (*δ*=128) on human data and from 11-14% (*δ*=32) to 3% (*δ*=128) on primate data, reflecting EB-TP’s faster per-segment re-alignment rather than slower decompression.

## 5 Discussion

Adaptive tracepoints address the limitations of fixed-length alignment compression by adapting segment boundaries to local sequence complexity. On simulated 100 Kb alignments, our method achieves 10.5-13.7× better compression than fixed-length encoding, while guaranteeing that reconstructed alignments maintain identical or improved scores. Diagonal-bounded tracepoints (DB-TP) consistently achieve the best compression because substitutions, which dominate genomic divergence, do not cause diagonal drift, yielding fewer and larger segments. Edit-bounded tracepoints (EB-TP) offer a tunable compression-cost trade-off: at *δ*=32, EB-TP provides moderate compression with substantially lower reconstruction cost; at *δ*=128, EB-TP approaches DB-TP compression (0.025× vs 0.025× on human data) while remaining 2-18× faster and requiring 4-13× less peak memory.

Reconstruction frequently improves upon input alignment scores. On the primate pangenome, 68.34%-80.07% of EB-TP (*δ*=32-128) and 75.66% of DB-TP reconstructions achieved better scores; on the human pangenome, 0.15%-3.15% (EB-TP) and 0.54% (DB-TP) improved, with zero degradations in all cases. This reflects the suboptimality of heuristic input alignments rather than a property of the encoding itself: tracepoint reconstruction applies exact WFA alignment within each segment, recovering optimal paths that heuristic aligners may miss, particularly for divergent sequences. By treating indels as atomic units, we further ensure biological fidelity: by never splitting indels across segment boundaries, our method preserves the interpretation of insertions and deletions of any size, a property that fixed-length approaches do not guarantee.

The integration with the TPA binary format achieves 23-139× compression on real pangenome data with indexed random access to individual alignment records. However, reconstruction cost remains a practical consideration: DB-TP’s larger segments require more resources for WFA alignment, with peak memory reaching 248 GiB for primate alignments. In many settings, EB-TP at higher *δ* offers a practical alternative, achieving near-equivalent compression with far lower reconstruction cost (Table 2). For workflows where decompression speed is important, BGZIP remains competitive at the cost of 7-39× more storage. The open-source implementation enables immediate integration into genomic pipelines managing large-scale pangenome comparisons. Although the format already supports edit-distance, gap-affine, and dual-affine gap scoring models, reconstruction currently recomputes the optimal wavefront band for each segment from the stored scoring parameters and segment dimensions. Future work will explore storing precomputed band bounds per segment to further accelerate CIGAR reconstruction, trading a small increase in file size for faster decompression. IMPG (Guarracino *et al*., 2025) already supports TPA files for querying compressed alignments for implicit pangenomics. In approximate mode, coordinate liftover and interval queries operate directly on the tracepoint metadata without sequence I/O or CIGAR reconstruction; in exact mode, only segments overlapping the queried range are reconstructed, avoiding full CIGAR recovery.

In a broader sense, tracepoint-based encoding presents a compact, structured summary of an alignment, analogous to a sketch of very large sequence data. Rather than storing the full alignment trace, tracepoints retain a sparse set of checkpoints that capture the global structure of the alignment while deferring fine-grained detail to on-demand reconstruction. This enables high-level operations on large alignments, such as indexing, filtering, and random access, without materializing the full CIGAR. In this respect, tracepoints expose alignment information at multiple resolutions, playing a role similar to sequence sketches in large-scale genomics, but with the additional guarantee that the original alignment can be reconstructed optimally when needed.

Overall, adaptive tracepoint encoding enables compression of large-scale alignments while preserving exact reconstructability and biological interpretability. By adapting tracepoint placement to local alignment complexity and supporting multiple scoring models, our approach provides a practical representation for storing and processing pangenome-scale alignment data. We expect this work to facilitate the development of more scalable alignment storage formats and analysis tools in future work.

## Funding

A.G., P.P., and E.G. are funded by NSF PPoSS Award #2118709; NIH R01HG013618; NIH U01HG013760; NIH U01DA057530; NIH U41HG010972; NIH R01HG013017; and the Tennessee Center for Integrative and Translational Genomics. S.M. is supported by the Spanish Ministry of Science and Innovation MCIN/AEI/10.13039/501100011033 (contracts PID2023-146193OB-I00 and PID2023-146511NB-I00), the Spanish Ministry, Ministerio para la Transformación Digital y de la Función Pública, and the European Union – NextGenerationEU through the Càtedra Chip UPC project (grant number TSI-069100-2023-0015).

## Data Availability

HPRCv2 assemblies can be downloaded from https://github.com/human-pangenomics/hprc_intermediate_assembly/tree/main/data_tables. T2T ape genome assemblies can be found in Yoo *et al*. (2025). Simulated sequences, and simulated and real alignments in TPA format are available at https://doi.org/10.5281/zenodo.18663011. Analysis code is available at https://github.com/chris7716/tracepoints-paper.

## Conflict of Interest

None declared.

## Notes

https://github.com/AndreaGuarracino/tracepoints

https://github.com/AndreaGuarracino/tpa

https://github.com/AndreaGuarracino/cigzip

https://github.com/chris7716/tracepoints-paper

https://doi.org/10.5281/zenodo.18663011

### Competing Interest Statement

The authors have declared no competing interest.

## References

Amarasinghe, S. L., Su, S., Dong, X., Zappia, L., Ritchie, M. E., and Gouil, Q. (2020). Opportunities and challenges in long-read sequencing data analysis. Genome Biology, 21(1).

Bonfield, J. K., Marshall, J., Danecek, P., Li, H., Ohan, V., Whitwham, A., Keane, T., and Davies, R. M. (2021). Htslib: C library for reading/writing high-throughput sequencing data. GigaScience, 10(2).

Bonfield, J. K., Marshall, J., Danecek, P., Li, H., Ohan, V., Whitwham, A., Keane, T., and Davies, R. M. (2022). CRAM 3.1: advances in the CRAM file format. Bioinformatics, 38(6), 1497–1503.

Chao, J., Tang, F., and Xu, L. (2022). Developments in algorithms for sequence alignment: A review. Biomolecules, 12(4), 546.

Gotoh, O. (1982). An improved algorithm for matching biological sequences. Journal of Molecular Biology, 162(3), 705–708.

Greenfield, D., Wittorff, V., and Hultner, M. (2019). The importance of data compression in the field of genomics. IEEE Pulse, 10(2), 20–23.

Guarracino, A., Mwaniki, N., Marco-Sola, S., and Garrison, E. (2021). wfmash: whole-chromosome pairwise alignment using the hierarchical wavefront algorithm.

Guarracino, A., Kille, B., and Garrison, E. (2025). impg: implicit pangenome graph.

Li, H., Handsaker, B., Wysoker, A., Fennell, T., Ruan, J., Homer, N., Marth, G., Abecasis, G., and Durbin, R. (2009). The sequence alignment/map format and samtools. Bioinformatics, 25(16), 2078–2079.

Marco-Sola, S., Moure, J. C., Moreto, M., and Espinosa, A. (2020). Fast gap-affine pairwise alignment using the wavefront algorithm. Bioinformatics, 37(4), 456–463.

Marco-Sola, S., Eizenga, J. M., Guarracino, A., Paten, B., Garrison, E., and Moreto, M. (2023). Optimal gap-affine alignment in o(s) space. Bioinformatics, 39(2).

Myers, E. W. (1986). An O(ND) difference algorithm and its variations. Algorithmica, 1(1-4), 251–266.

Myers, G. (2014). Efficient local alignment discovery amongst noisy long reads. In Algorithms in Bioinformatics (WABI 2014), volume 8701 of Lecture Notes in Computer Science, pages 52–67. Springer Berlin Heidelberg.

Myers, G., Durbin, R., and Zhou, C. (2025). Fastga: fast genome alignment. Bioinformatics Advances, 5(1), vbaf238.

Needleman, S. B. and Wunsch, C. D. (1970). A general method applicable to the search for similarities in the amino acid sequence of two proteins. Journal of molecular biology, 48(3), 443–453.

Smith, T. and Waterman, M. (1981). Identification of common molecular subsequences. Journal of Molecular Biology, 147(1), 195–197.

Stephens, Z. D., Lee, S. Y., Faghri, F., Campbell, R. H., Zhai, C., Efron, M. J., Iyer, R., Schatz, M. C., Sinha, S., and Robinson, G. E. (2015). Big data: Astronomical or genomical? PLOS Biology, 13(7), e1002195.

Yoo, D., Rhie, A., Hebbar, P., Antonacci, F., Logsdon, G. A., Solar, S. J., Antipov, D., Pickett, B. D., Safonova, Y., Montinaro, F., Luo, Y., Malukiewicz, J., Storer, J. M., Lin, J., Sequeira, A. N., Mangan, R. J., Hickey, G., Monfort Anez, G., Balachandran, P., Bankevich, A., Beck, C. R., Biddanda, A., Borchers, M., Bouffard, G. G., Brannan, E., Brooks, S. Y., Carbone, L., Carrel, L., Chan, A. P., Crawford, J., Diekhans, M., Engelbrecht, E., Feschotte, C., Formenti, G., Garcia, G. H., de Gennaro, L., Gilbert, D., Green, R. E., Guarracino, A., Gupta, I., Haddad, D., Han, J., Harris, R. S., Hartley, G. A., Harvey, W. T., Hiller, M., Hoekzema, K., Houck, M. L., Jeong, H., Kamali, K., Kellis, M., Kille, B., Lee, C., Lee, Y., Lees, W., Lewis, A. P., Li, Q., Loftus, M., Loh, Y. H. E., Loucks, H., Ma, J., Mao, Y., Martinez, J. F. I., Masterson, P., McCoy, R. C., McGrath, B., McKinney, S., Meyer, B. S., Miga, K. H., Mohanty, S. K., Munson, K. M., Pal, K., Pennell, M., Pevzner, P. A., Porubsky, D., Potapova, T., Ringeling, F. R., Rocha, J. L., Ryder, O. A., Sacco, S., Saha, S., Sasaki, T., Schatz, M. C., Schork, N. J., Shanks, C., Smeds, L., Son, D. R., Steiner, C., Sweeten, A. P., Tassia, M. G., Thibaud-Nissen, F., Torres-González, E., Trivedi, M., Wei, W., Wertz, J., Yang, M., Zhang, P., Zhang, S., Zhang, Y., Zhang, Z., Zhao, S. A., Zhu, Y., Jarvis, E. D., Gerton, J. L., Rivas-González, I., Paten, B., Szpiech, Z. A., Huber, C. D., Lenz, T. L., Konkel, M. K., Yi, S. V., Canzar, S., Watson, C. T., Sudmant, P. H., Molloy, E., Garrison, E., Lowe, C. B., Ventura, M., O’Neill, R. J., Koren, S., Makova, K. D., Phillippy, A. M., and Eichler, E. E. (2025). Complete sequencing of ape genomes. Nature, 641(8062), 401–418.

