## Supplementary Material for "Adaptive Tracepoints for Pangenome Alignment Compression"

Hasitha Kaushan[1], Santiago Marco-Sola[2,3], Erik Garrison[4], Pjotr Prins[4], and Andrea Guarracino[4,5,*]

[1]Department of Electrical Engineering, University of Moratuwa, Bandaranayake Mawatha, Moratuwa 10400, Sri Lanka
[2]Barcelona Supercomputing Center, Universitat Politècnica de Catalunya, Barcelona, Spain
[3]Department of Computer Science, Universitat Politècnica de Catalunya, Spain
[4]Department of Genetics, Genomics and Informatics, University of Tennessee Health Science Center, Memphis, TN 38163, USA
[5]Bioinnovation and Genome Sciences, The Translational Genomics Research Institute (TGen), Phoenix, AZ 85004, USA

**Summary**

This supplementary material complements the main manuscript by presenting formal proofs of reconstruction optimality under edit-distance and affine-gap scoring models, banding lemmas for accelerated reconstruction, TPA format details, and supplementary figures for simulated and real pangenome experiments.

### 1 Optimal Edit-Distance Alignment Reconstruction using Tracepoints

Let $q_{1,\cdots,n}$ and $t_{1,\cdots,m}$ be two sequences, and let $A^\star$ denote an optimal edit-distance alignment path in the standard edit graph. Let $T = \{(x_0, y_0), \ldots, (x_N, y_N)\}$ be a sequence of tracepoints sampled from $A^\star$, ordered such that $0 = x_0 < x_1 < \cdots < x_N = n$ and $0 = y_0 < y_1 < \cdots < y_N = m$ and each $(x_i, y_i)$ lies on $A^\star$. For every $i \in \{0, \ldots, N-1\}$, let $A_i$ be an optimal alignment between the substrings $q_{x_i+1,\cdots,x_{i+1}}$ and $t_{y_i+1,\cdots,y_{i+1}}$, constrained to start at $(x_i, y_i)$ and end at $(x_{i+1}, y_{i+1})$.

**Lemma 1.1** (Edit-distance optimality via reconstruction between tracepoints)**.** *Let $A = A_0 \circ A_1 \circ \cdots \circ A_{N-1}$ be the concatenation of the optimal edit-distance subalignments between tracepoints. Then, $A$ is an optimal alignment between $q$ and $t$ under the edit-distance model (but not necessarily identical to $A^\star$).*

*Proof.* The alignment problem can be formulated as a minimum-cost path in the acyclic edit graph whose vertices are grid points $(i, j)$, with additive edge costs for matches, mismatches, and gap penalties. Because the total cost of any alignment path is additive, an optimal path $A^\star$ from $(0, 0)$ to $(n, m)$ can be decomposed into subpaths (subproblems) between consecutive tracepoints $(x_i, y_i)$ and $(x_{i+1}, y_{i+1})$.

Assume by contradiction that there exists a strictly cheaper subpath $\tilde{A}_i$ from $(x_i, y_i)$ to $(x_{i+1}, y_{i+1})$ than the corresponding segment $A_i^\star$ of $A^\star$. Replacing $A_i^\star$ with $\tilde{A}_i$ in $A^\star$ yields a new global path $\tilde{A}$ with a strictly smaller total cost, contradicting the optimality of $A^\star$. Therefore, each segment $A_i^\star$ must already be optimal between its corresponding tracepoints.

Since each $A_i$ is by definition an optimal alignment between the same endpoints, concatenating them yields a global alignment $A = A_0 \circ A_1 \circ \cdots \circ A_{N-1}$ of equal minimal cost to $A^\star$. Hence, $A$ is an optimal alignment between $q_{1,\cdots,n}$ and $t_{1,\cdots,m}$.

Note that, because optimal edit-distance alignments are not necessarily unique, the reconstructed alignment is guaranteed to be optimal in total cost but may differ from $A^\star$ in the specific sequence of edit operations, since multiple distinct paths can achieve the same minimum cost between the same endpoints. □

**Corollary 1.1.1.** *Let $T$ be any sequence of tracepoints extracted from an alignment path $A$ (not necessarily optimal). If each subalignment between consecutive tracepoints is recomputed optimally, then the reconstructed alignment $\hat{A}$ has total cost less than or equal to that of $A$. In particular, if $A$ is not optimal, the reconstructed alignment cannot be worse (i.e., cannot have a higher edit distance) and may strictly improve it.*

### 2 Optimal Affine-Gap Alignment Reconstruction using Tracepoints

In the affine-gap model, a gap of length $\ell$ is penalized by a cost of the form $g_o + \ell \cdot g_e$, where $g_o$ and $g_e$ denote the gap opening and gap extension penalties, respectively. This scoring function better captures the biological reality of insertions and deletions occurring in contiguous runs, but it also breaks the one-to-one correspondence between diagonal motion and edit count that underlies the unit-cost analysis.

Under an affine-gap scoring model, if a tracepoint splits a contiguous insertion or deletion into two disjoint parts, then independently reconstructing the two resulting subalignments is not guaranteed to yield the same total score as the globally optimal alignment. Note that the cost of a gap depends on whether it is represented as a single contiguous segment or fragmented into multiple segments, since each fragment incurs its own gap-opening penalty. Suppose that, in the globally optimal alignment, a gap of total length $\ell$ crosses a tracepoint and is therefore split into two parts of lengths $\ell_1$ and $\ell_2$, with $\ell_1 + \ell_2 = \ell$. The true global cost of this gap is $g_o + \ell g_e$. However, if the two subalignments are reconstructed independently, each part is treated as a separate gap and could incur its own opening penalty, yielding a combined cost of $(g_o + \ell_1 g_e) + (g_o + \ell_2 g_e) = 2g_o + \ell g_e$ which is strictly larger than the global cost.

**Lemma 2.1** (Affine-safe tracepoint reconstruction under atomic gaps)**.** *Consider an optimal alignment under an affine-gap scoring model, and let $T = \{(x_0, y_0), \ldots, (x_N, y_N)\}$ be a sequence of tracepoints sampled along this alignment such that no tracepoint lies strictly inside a gap. For each $i \in \{0, \ldots, N-1\}$, let $A_i$ be an optimal affine-gap alignment between the substrings $q_{x_i,\cdots,x_{i+1}-1}$ and $t_{y_i,\cdots,y_{i+1}-1}$ constrained to start at $(x_i, y_i)$ and end at $(x_{i+1}, y_{i+1})$. Then, the concatenation $A = A_0 \circ A_1 \circ \cdots \circ A_{N-1}$ is a globally optimal alignment under the affine-gap scoring model.*

*Proof.* By construction, the original optimal alignment induces a path that passes through all tracepoints in $T$, and, by hypothesis, no tracepoint falls strictly inside a gap. Hence, each affine gap segment in the optimal alignment is fully contained within a single interval between two consecutive tracepoints and is not split across subalignments.

Therefore, the conditions of Lemma 1.1 apply to the affine-gap edit graph: each $A_i$ is, by definition, an optimal affine-gap alignment between its corresponding pair of tracepoints, and the affine-gap cost is additive across these unsplit segments. It follows that concatenating the subalignments yields a global alignment $A = A_0 \circ A_1 \circ \cdots \circ A_{N-1}$ with the same minimal total cost as the original globally optimal alignment under the affine-gap scoring model. □

### 3 Accelerating Alignment Reconstruction by Exploiting Local Edit-Bounds

By knowing the number of edit operations (or alignment score) in each subalignment between tracepoints, we can explicitly bound the width of the diagonal band required to contain the optimal path for that segment. As a result, the reconstruction alignment algorithm no longer needs to explore the whole region, but only a narrow band whose width is proportional to the local edit bound.

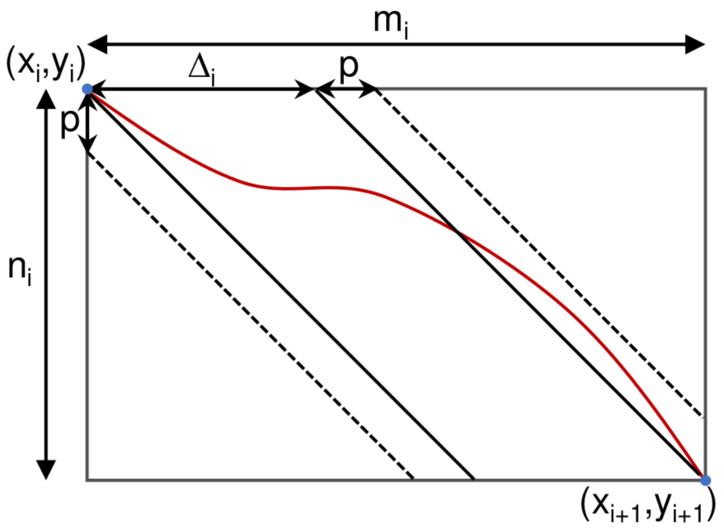


Figure 1: Diagram of a subalignment between two tracepoints. Reconstruction using local edit-bounds via banded alignment.

Given two tracepoints $(x_i, y_i)$ and $(x_{i+1}, y_{i+1})$, we define $n_i = y_{i+1} - y_i$, $m_i = x_{i+1} - x_i$, as the lengths of the corresponding substrings in $q$ and $t$, respectively. Let $e_i$ denote the number of edit operations (i.e., insertions, deletions, and mismatches) in the optimal subalignment between $q_{y_i,\cdots,y_{i+1}-1}$ and $t_{x_i,\cdots,x_{i+1}-1}$. We further define the diagonal drift as $\Delta_i = |m_i - n_i| = |\#\text{ins}_i - \#\text{del}_i|$.

**Lemma 3.1** (Diagonal band bound under edit-distance). *The optimal alignment path between these two consecutive tracepoints $(x_i, y_i)$ and $(x_{i+1}, y_{i+1})$ is contained within a band of diagonals $k$, where $-p \leq k \leq \Delta_i + p$ and $p = (e_i - \Delta_i)/2$.*

*Proof.* Without loss of generality, let's suppose that $n_i < m_i$. As Figure 1 shows, to go from $(x_i, y_i)$ to $(x_{i+1}, y_{i+1})$ the path must contain a total of $\Delta_i = |m_i - n_i|$ insertions. This means that at least $\Delta_i$ of the $e_i$ edit operations are mandatorily to reach $(x_{i+1}, y_{i+1})$. The remaining $e_i - \Delta_i$ edit operations may allow the path to drift away from the central diagonal band $0 \leq k \leq \Delta_i$. However, any such deviation must be eventually compensated to return to the required final offset. Therefore, the maximum number of extra diagonal steps that can be spent drifting on either side is $p = \frac{e_i - \Delta_i}{2}$. It follows that the diagonal offset $k$ along the entire optimal subalignment path is bounded by $-p \le k \le \Delta_i + p$, which proves the claim. □

Similarly, consider the optimal affine-gap subalignment between two consecutive tracepoints $(x_i, y_i)$ and $(x_{i+1}, y_{i+1})$ with total alignment cost $s$. Let the mandatory drift cost under the affine-gap model be $\Delta g_e$.

**Lemma 3.2** (Diagonal band bound under affine-gap)**.** *The optimal alignment path between* $(x_i, y_i)$ *and* $(x_{i+1}, y_{i+1})$ *is contained within the band of diagonals* $k$ *satisfying*

$$-p_{\text{gap}} \le k \le \Delta_i + p_{\text{gap}}, \qquad \textit{where} \qquad p_{\text{gap}} = \max\left\{\frac{s - 2g_o - \Delta g_e}{2g_e}, 0\right\}.$$

*Proof.* Similar to the proof of Lemma 3.1. The diagonal drift $\Delta_i$ is fixed by the endpoints. Any feasible alignment must therefore perform at least $\Delta_i$ gap extensions in total, and hence incur at least $\Delta g_e$ cost. The remaining budget $(s - \Delta g_e)$ can only be spent on additional deviations (detours) from the central diagonal strip $0 \le k \le \Delta_i$. Any unit deviation beyond this strip must be compensated later to reach the required endpoint diagonal, which costs at least two gap extensions: one to move away and one to return. Moreover, under affine penalties, such a detour requires indels in both directions (to drift away and to return), which induces two gap segments and therefore requires paying two gap opening penalties in total. Thus, at least $2g_o$ cost units must be reserved for gap openings when accounting for the maximum detour. Consequently, the maximum number of extra diagonal steps that can be spent drifting on either side is bounded by $p_{\text{gap}} = \max\{(s - 2g_o - \Delta g_e)/2g_e, 0\}$. Combining both sides yields $-p_{\text{gap}} \le k \le \Delta_i + p_{\text{gap}}$, as claimed. □

### 4 TPA Encoding Details

TPA decomposes each alignment record into two independent streams that are encoded separately. The content of these streams depends on the tracepoint type:

- **EB-TP and DB-TP** store $(a_{\text{len}}, b_{\text{len}})$ pairs representing query and target advances per segment. The first stream carries $a_{\text{len}}$ values and the second carries $b_{\text{len}}$ values.
- **FL-TP** stores a per-segment edit distance and a target advance $(b_{\text{len}})$. The first stream carries edit distances and the second carries $b_{\text{len}}$ values.

Table 1 summarizes the encoding strategy selected for each stream under each tracepoint type.

Table 1: TPA encoding strategies per tracepoint type and stream.

| TP type | First stream | Second stream |
|---|---|---|
| EB-TP | Rice coding | Delta + zigzag varint |
| DB-TP | Raw varint | Delta + zigzag varint |
| FL-TP | Rice coding | Rice coding |

The encoding strategies are:

- **Rice coding**: each value $v$ is decomposed as $v = q \cdot 2^k + r$, where the quotient $q$ is unary-coded ($q$ one-bits followed by a zero-bit) and the remainder $r$ is stored in $k$ fixed bits. The parameter $k$ is auto-tuned per record by testing $k \in \{0, \ldots, 16\}$ and choosing the value that minimizes the total encoded size. This adds a 1-byte header per record to store $k$.

- **Raw varint** (LEB128): each value is encoded using 7 data bits per byte with one continuation bit; values below 128 fit in a single byte, below 16,384 in two bytes, and so on. No per-record overhead.

- **Delta + zigzag varint**: each second-stream value is encoded as the signed residual $b_{\text{len}} - a_{\text{len}}$, mapped to an unsigned integer via zigzag encoding ($v \mapsto (v \ll 1) \oplus (v \gg 63)$) so that small absolute differences produce small unsigned values, then stored as a varint. No per-record overhead.

**Second stream.** For EB-TP and DB-TP, query and target advances are highly correlated (Pearson $r > 0.9$ on both intra- and cross-species alignments), so the second stream is encoded as a delta from the first: the residual $b_{\text{len}} - a_{\text{len}}$ is typically small and most values fit in a single byte as zigzag variable-length integers. For FL-TP, the two streams measure different quantities (edit distance vs. target advance) and are effectively uncorrelated ($r < 0.1$), so delta encoding between them is not applied. Instead, each stream is independently Rice-coded, exploiting the right-skewed distribution of both value types.

**First stream.** Under EB-TP, edit-distance segmentation produces right-skewed segment-length distributions; Rice coding exploits this geometric-like shape at bit-level precision. Under DB-TP, diagonal-distance segmentation produces fewer but larger segments; Rice coding's per-record overhead outweighs its bit-level savings, making raw variable-length integers more compact. Under FL-TP, per-segment edit distances follow a right-skewed distribution; Rice coding provides efficient bit-level encoding for these values with minimal per-record overhead.

**Optional block compression.** An optional per-record compression layer (zstd) can be applied when records contain enough tracepoints to amortize the compressor's overhead. In practice, the metric-specific encodings already capture most of the redundancy, and the additional compression layer provides marginal benefit across the datasets tested.

### 5 Supplementary Figures

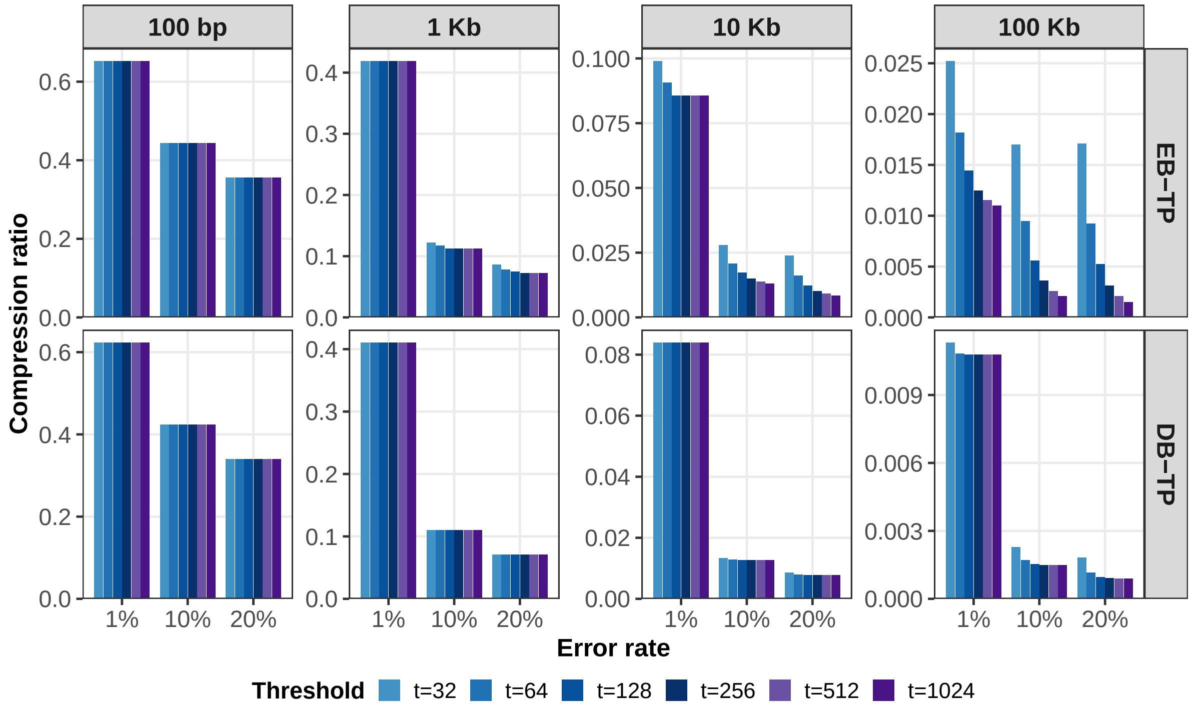


Figure 2: Effect of threshold parameter on compression ratio for EB-TP and DB-TP on simulated data ($N = 10{,}000$ pairs per condition). Each panel shows a method–length combination. Bars represent threshold values $t \in \{32, 64, 128, 256, 512, 1024\}$ (where $t = \delta$ for EB-TP and $t = b$ for DB-TP) at error rates of 1%, 10%, and 20%. Lower ratios indicate better compression.

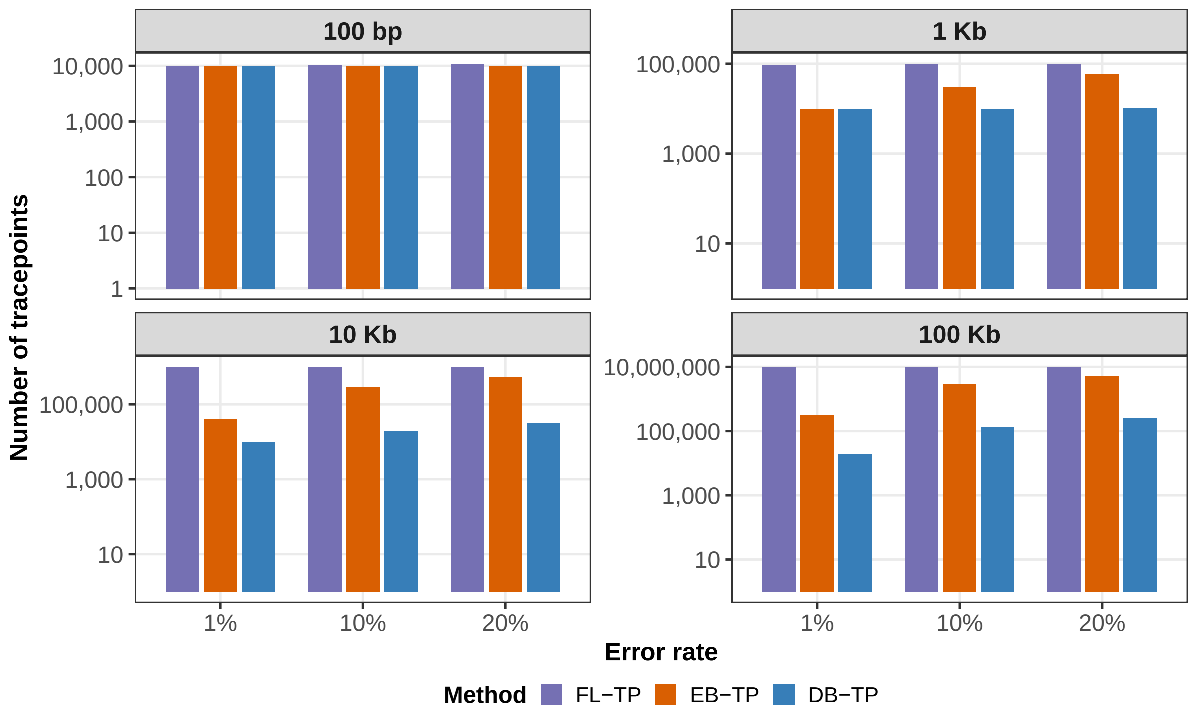


Figure 3: Number of tracepoints per method on simulated data. Panels: sequence lengths (100 bp–100 Kb). FL-TP ($l = 100$) produces a fixed count regardless of divergence, while EB-TP ($\delta = 32$) and DB-TP ($b = 32$) scale with alignment complexity, generating up to $\sim 500\times$ fewer tracepoints at low divergence.

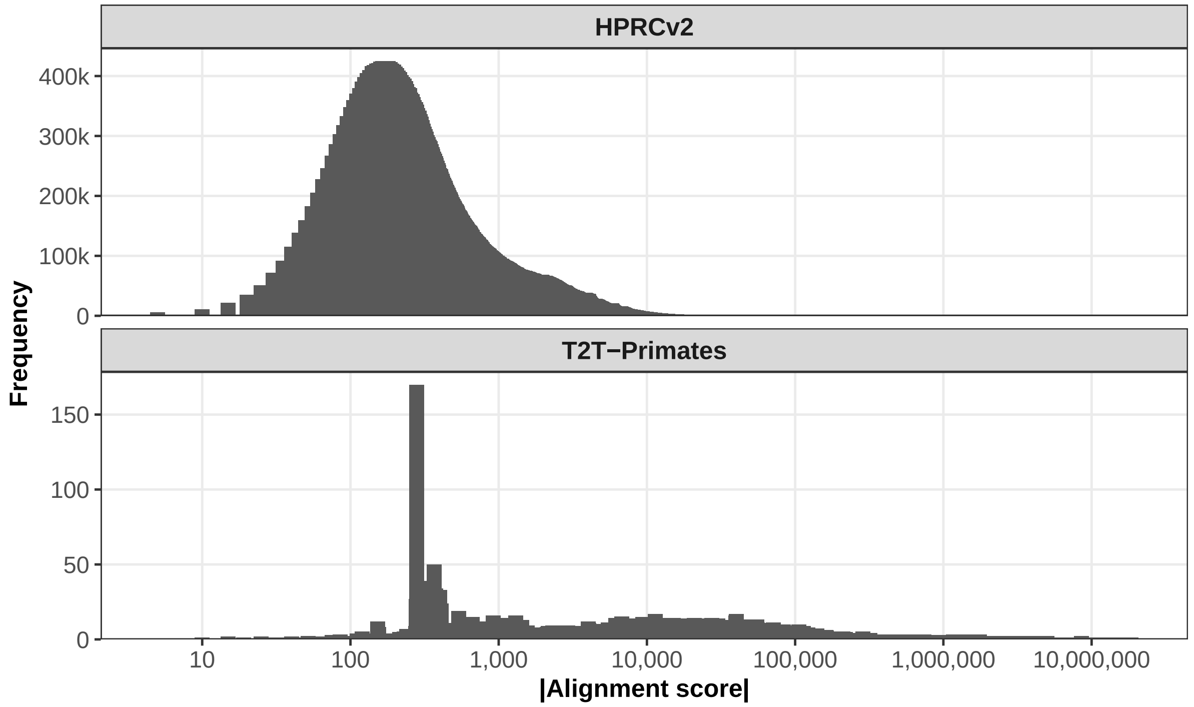


Figure 4: Alignment score distributions for the two pangenomes. Top: human pangenome (HPRCv2, 389M alignments, 466 haplotypes). Bottom: primate pangenome (T2T apes, 566K inter-species alignments, 20 haplotypes, 7 ape species). The primate pangenome shows a broader distribution reflecting higher inter-species divergence.

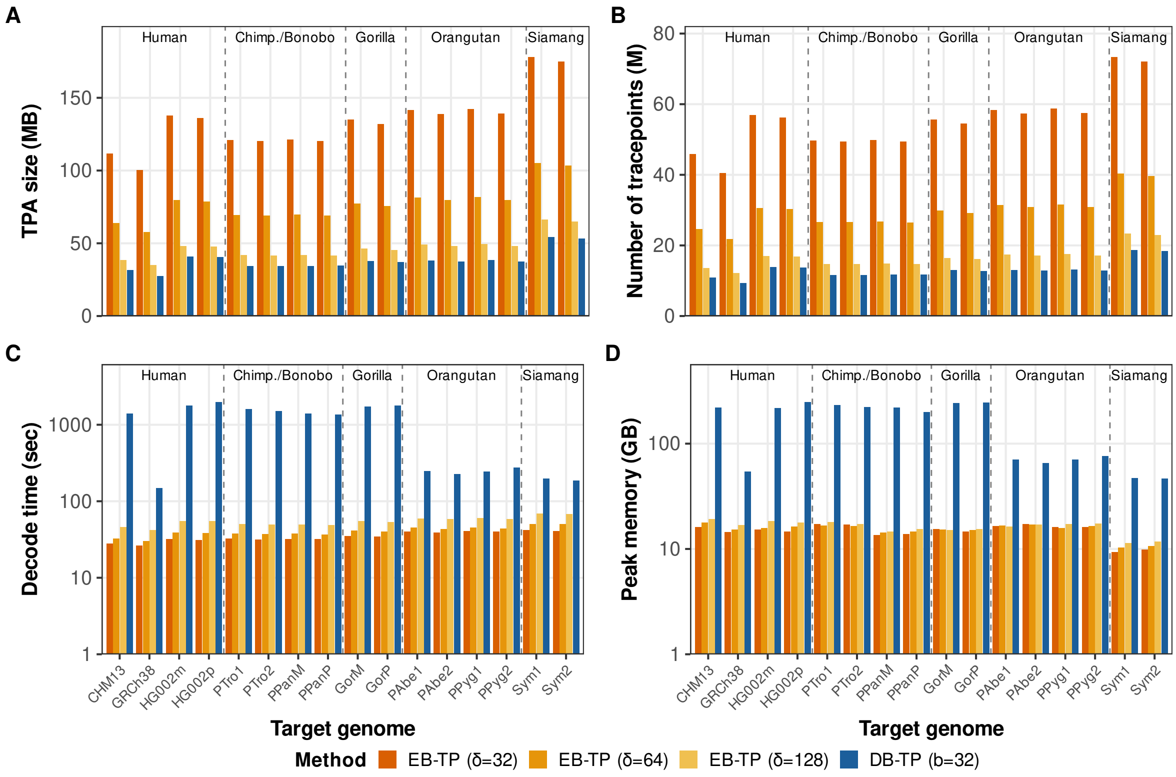


Figure 5: Reconstruction performance on the primate pangenome by target genome. (A) TPA file size. (B) Number of tracepoints. (C) Decoding time. (D) Peak memory. Targets are grouped by closely related species. Three EB-TP thresholds ($\delta = 32, 64, 128$) are compared with DB-TP ($b = 32$). Increasing $\delta$ reduces file size and tracepoint count but increases decoding time. DB-TP ($b = 32$) achieves the smallest files and fewest tracepoints, at the cost of substantially higher decoding time and memory, particularly for more divergent species pairs.
